# Evaluation of the Diuretic Activities of 80% Methanol Leaf Extract and Solvent Fractions of *Rumex nepalensis* in Mice

**DOI:** 10.1101/2023.11.10.566525

**Authors:** Fasika Argaw Tafesse, Assefa Belay Asrie, Tafere Mulaw Belete

## Abstract

**Background:** The leaf of *Rumex nepalensis* has historically been employed to treat urinary retention and as a diuretic. Despite these assertions, there has been very little research addressing the diuretic effect of the leaves of *R. nepalensis*. Therefore, this study was done to examine the diuretic properties of R. nepalensis leaves.

**Methods:** Cold maceration with 80% methanol was used to extract the coarsely powdered leaves of *R. nepalensis*. The extract was separated using increasing polarity solvents, beginning with n-hexane, ethyl acetate, and water. Mice were split into sections to test the plant’s diuretic properties. The negative control groups were given either distilled water or 2% tween 80; the positive control groups were given furosemide (10 mg/kg); and the test groups were given the 80% methanol extract and solvent fractions orally at dosages of 100, 200, and 400 mg/kg. The urine volume was determined, and urine analysis was performed on each extract.

**Results:** At dosage levels of 200 mg/kg and 400 mg/kg, the 80% methanol extract, ethyl acetate fraction, and aqueous fraction all produced substantial diuresis (p<0.001) as compared to the negative control. Similarly, mice given the 80% methanol extract, ethyl acetate fraction, and aqueous fraction demonstrated substantial natriuresis (p<0.001) and kaliuresis (p<0.001) at dosages of 200 mg/kg and 400 mg/kg, respectively, in comparison to the negative control.

**Conclusion:** The diuretic activity of *R. nepalensis* was significantly seen in the 80% methanol extract, ethyl acetate, and aqueous fractions, which corroborated the plant’s traditional use.

## Introduction

Diuretics are drugs which enhance urine flow by causing the kidneys to remove electrolytes and water from the body.(1) They are among the most commonly prescribed medications, and are used to establish a negative fluid and salt balance in a number of clinical disorders.(2) However, their use has been attributed to a range of problems that are predicted given their modes of action, including hypokalemia, hyponatraemia, metabolic acidosis, glucose intolerance, and hypomagnesaemia.(3) Furthermore, the global prevalence of diuretic resistance is projected to be 21% among heart failure patients. (4) Despite the variations in approaches taken to reduce these unpleasant consequences, there is an urgent need for newer and better diuretics with unique modes of action, enhanced efficacy, ease of accessibility and cost, and tolerable side effect profiles.

Over fifty percent of the medications that are in the Western pharmacopoeia are derived from herbs or developed from adaptations of substances previously recognized in plants, making plant substances the cornerstone of modern medicine.(5) *R. nepalensis* (often termed as Nepal Dock) belongs to the Polygonaceae family. The genus includes roughly 250 species of plants that are scattered globally.(6) Japan, India, Indonesia, Myanmar, Nigeria, Ethiopia, and the Republic of South Africa are among the countries where the plant is found to thrive.(7) The vernacular names of *R. nepalensis* are Yewusha tult in Amharic (8), Shuultii/Tulet in Oromiffaa, Oromia Regional State, Ethiopia.(9) The plant grows in all seasons of the year, and has broad roots, upright stems, with basal leaves up to 10 cm long.(10)

Extracts of *R. nepalensis* are claimed to exhibit anti-ulcer (11), abortifacient (8), antioxidant and anti-inflammatory (12), hypoglycemic and analgesic (13), antiplasmodial and antimicrobial (14), and antimalarial activities.(15) These investigations have led to the separation of secondary phytochemicals from the plant such as alkaloids, flavonoids, phenolic compounds, and glycosides. (8, 11) Furthermore, The root of *R. nepalensis* is commonly used for urine retention in Ethiopia, while its leaves are used as a diuretic in two parts of Pakistan. (9, 16, 17) As these applications are not examined scientifically, The current study was carried out to assess the diuretic effects of *R. nepalensis* (Polygonacea) leaves 80% methanol extract and solvent fractions in mice.

## Material and methods

### Chemicals and reagents

Distilled water (EPHARM Ethiopia), absolute methanol (Follium Pharmaceuticals, Ethiopia), ethyl acetate (Sisco research laboratories Pvt. Ltd, India), n-hexane (Loba Chemie Pvt. Ltd. India), normal saline (EPHARM Ethiopia), furosemide 40 mg tablet (Salutas Pharma GmBH, Germany), tween 80 (Care laboratories and Medical supplies, India) were purchased from their respective suppliers. All of the chemicals and reagents utilized were of analytical grade.

### Instruments, Apparatuses and Supplies

Metabolic cage, oral gavage, urine collector, digital weighing balance, lyophilizer, rotary evaporator (YAMATO rotary evaporator RE301, Japan), deep freezer, digital pH meter (HANNA instruments, Italy), vacuum pump (AB-288-I, India), mortar and pestle, separatory funnel, beaker, Erlenmeyer conical flask, filter paper (Whatman No. 1, GE Bio-Sciences, Thailand), examination glove (ASAP International SDN BHD, Malaysia), and gauze were employed.

### Plant material

The leaves of *R. nepalensis* were gathered from Atse-Tewodros Campus, University of Gondar, Gondar Town, Amhara region at a longitude and latitude of 37^0^ 26’ 3” E, 12^0^ 35’ 20” N, respectively and an elevation of 2177 meters. A botanist from the Department of Biology, College of Natural and Computational Sciences, University of Gondar, Gondar, Ethiopia, identified the plant samples, and a voucher specimen number of 01/FA/2022 was placed in the herbarium for future reference. The gathered material was cleaned with running water to eliminate pollutants and dust before drying in the shade for two weeks. The dry material was then roughly pulverized with a mortar and pestle.

### Experimental animals

We employed Swiss albino mice that were raised in the animal house of the Department of Pharmacology at the University of Gondar and weighed between 25 and 35 g. Six animals were housed in each polypropylene cage, and the environment was maintained at ambient temperature with 12-hour light and 12-hour dark cycles. They were given a typical pellet diet and unlimited access to water. One week before the commencement of the experiment, the animals became accustomed to the lab setting. All treatment and care of the animals during the experiment complied with the standards for the care and use of laboratory animals.(18)

### Preparation of hydromethanolic crude extract

Three Erlenmeyer conical flasks were used to macerate one kilogram of the R. nepalensis leaves with a total of 8000 mL of 80% methanol.(11) The flasks were wrapped in aluminum foil and left at room temperature for three days while being shaken now and then. Following three days, the extract was filtered first through a gauze and then through Whatman’s filter paper No1. The yield was then maximized by pressing and re-macerating the marc twice with the same solvent and under the same circumstances. A rotary evaporator with a temperature setting of 40^0^C and a rotation speed of 60 rpm was then used to concentrate the filtrates. A lyophilizer was used to freeze dry the concentrated aqueous solution after it had been frozen in a deep freezer during the previous night. The % yield of the extract was then determined, labelled, and stored in a refrigerator at a temperature of 4°C until use.

### Fractionation of crude hydromethanolic extract

The 80% crude methanol extract was fractionated further using a separatory funnel device and solvents of increasing polarity (n-hexane, ethyl acetate, and distilled water). Seventy grams of dried 80% crude methanol extract were suspended in 250 mL of distilled water and gently agitated to thoroughly dissolve. The solution was poured into a separating funnel. The mixture was then treated with an equal amount of n-hexane. After a quick shake, the liquid was given a few minutes to settle before being divided into two layers. Because n-hexane has a lower density than water, it was extracted from the upper phase. The same process, as described above, was used to obtain the whole hexane fraction. After collecting the hexane fraction, an equivalent amount (250 mL) of ethyl acetate was added to the residual aqueous part in the separating funnel, which was vigorously shaken. The process was performed again since the top layer, which was ethyl acetate, was separated from the aqueous component. The aqueous fraction was then collected. The n-hexane and ethyl acetate filtrates were then concentrated in a rotary evaporator at 40°C and 60 revolutions per minute. The aqueous fraction was freeze-dried using a lyophilizer after being frozen overnight in a deep freezer. Finally, the dried fraction yields were determined, placed into separate vials, and kept at -4°C.

### Grouping and dosing of animals

Mice were randomly divided into fifteen groups of six (n = 6). They were divided into two negative control groups, one standard group, and the others (groups IV through XV) as *R.nepalensis* treatment groups at random. The first negative control group received distilled water, whereas the second received 2% Tween 80. Furosemide 10 mg/kg was given to the standard group. The three different test dosages (100, 200, and 400 mg/kg) of *R.nepalensis* leaves 80% methanol extract were given to groups IV through VI. The remaining groups (Groups VII–XV) were given three independent test doses of n-hexane, ethyl acetate, and the aqueous fractions (100, 200, and 400 mg/kg).

### Acute oral toxicity test

Acute toxicity testing was carried out on nulliparous and non-pregnant female mice aged 8-12 weeks using the limit test approach outlined in the Organization for Economic Cooperation and Development Guideline No-425.(19) The LD_50_ of *R. nepalensis* crude extract was determined in this test using five female mice aged 8-12 weeks. All animals were fasted for four hours prior to and two hours after the extract was administered. One animal was given a limit dosage of 2000 mg/kg of the extract at first. Because the animal was shown to be safe after the first 24 hours, the extract was given to four other animals. The animals were then kept singly and examined for four hours at 30 minute intervals for a total of two weeks for any symptoms of toxicity, illness, or death.

### Determination of diuretic activity

The techniques employed by Wubshet Hailu and Ephrem Engidawork to assess diuretic action were utilized.(20) Prior to the experiment’s start, each mouse was housed in a separate metabolic cage for 24 hours to allow for adaptation. After that, the mice fasted for the whole night while having unrestricted access to water. To apply a consistent water and salt load, the animals received a pretreatment of physiological saline (0.9% NaCl) through oral administration at a dosage of 0.15 ml per 10 g of body weight.(21) Then, according to the grouping and dosage section, the treatment was administered to each animal orally through gavage. Then mice were promptly placed into metabolic cages (one mouse per cage) that are specifically designed to prevent the urine from mixing with feces. After that, urine was measured and collected for examination at the first, second, third, fourth, and fifth hours after the dose. It was then kept at - 20°C. Water and food were not provided to the animals throughout the urine collection process. The total volume of urine, Na+, K+, and Cl-concentrations, as well as the pH of the urine, were all assessed for every mouse. In order for comparison of the effects of the extracts with those of the vehicle and standard in terms of diuresis, the parameters of urine excretion, diuretic action, and diuretic activity were assessed. Calculated using **Formula 1**, total urine output was divided by total liquid consumed to get the urinary excretion independent of animal weight. The diuretic effect of a particular dosage of a drug was assessed by comparing the urine excretion in the test group to the urinary excretion in the control group (**Formula 2**). For comparison of the extract’s diuretic effect to that of the standard group’s reference drug (furosemide), a third measure known as diuretic activity was computed (**Formula 3**).

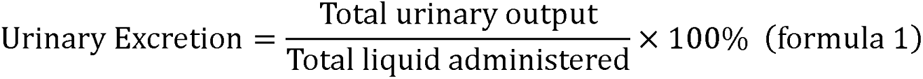

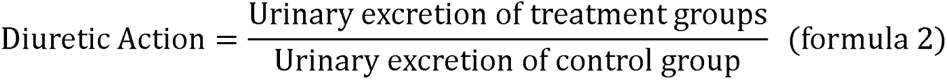

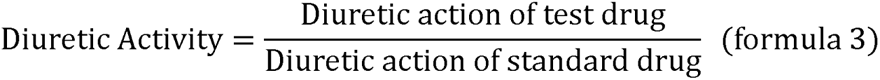

### Urine electrolyte analysis

Using an Ion Selective Electrode (ISE) analyzer (Beckman Coulter DxC 700 AU, USA), the concentrations of Na^+^, K^+^, and Cl^-^ in the urine were measured in all groups. Electrolyte concentration in urine was measured in mmol/L. The natriuretic index and carbonic anhydrase inhibition were assessed using the electrolyte ratios Na^+^/K^+^ and Cl^-^/K^+^+Na^+^, respectively. Additionally, the pH of urine samples was assessed using a digital pH meter. For the sake of ruling out their impact on urinary electrolyte concentration, the salt level of the crude extract and fractions was also measured using the same instrument.

### Phytochemical screening test

Using accepted techniques, the presence of secondary metabolites like alkaloids, anthraquinones, flavonoids, glycosides, phenols, saponins, phytosterols, tannins, and triterpenoids in the crude extract and solvent fractions of *R. nepalensis* leaves was qualitatively evaluated.(22)

### Data analysis

SPSS version 24 was used to analyze the data. The statistical significance of each set of data was determined using one-way analysis of variance (ANOVA), followed by Tuckey’s multiple comparison test. Mean plus or minus standard error of the mean (M±SEM) was used to describe the study’s findings. A P-value of <0.05 was used to determine if mean differences were statistically significant.

### Ethical clearance

The Department of Pharmacology, College of Medicine and Health Sciences, University of Gondar, granted ethical approval with the reference number SOP 4/57/2014.

## Results

### Percentage yield

With a percentage yield of 18.9%, 189 g dry powder was produced from the crude 80% methanol extract. (See **Table 1**).

**Table 1.**
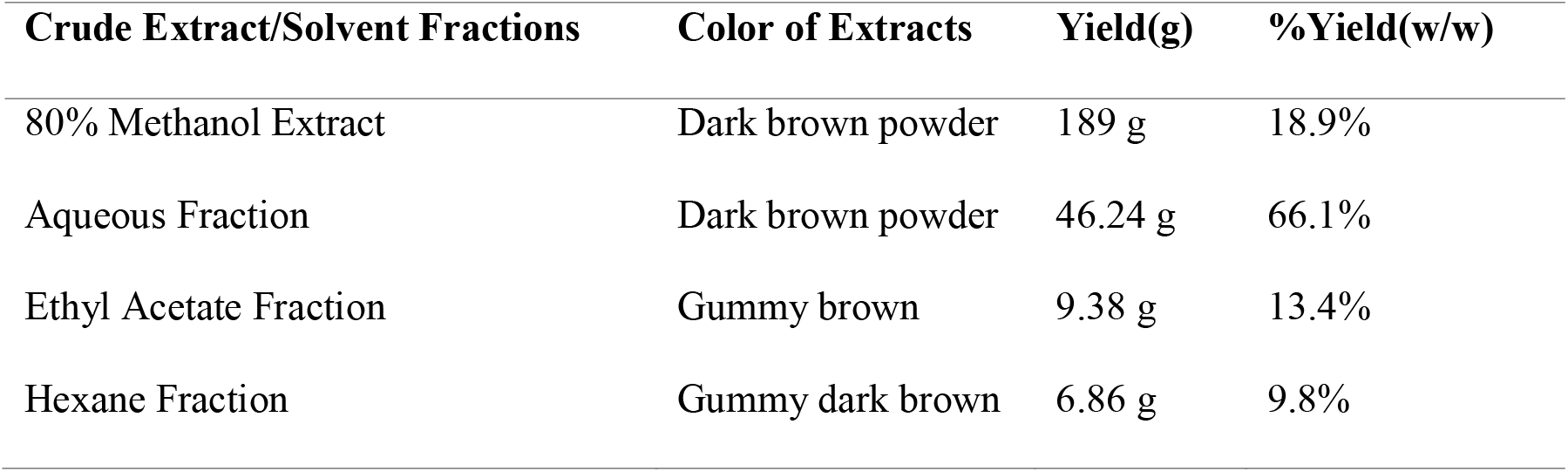
Amount, color, and percentage yields of the crude extract and fractions.

### Preliminary phytochemical screening

The presence of alkaloids, anthraquinones, flavonoids, glycosides, phenolic compounds, plant steroids, saponins, tannins, and terpenoids was revealed in an 80% methanol crude extract of *R. nepalensis* leaf (see **Table 2**).

**Table 2.** Preliminary phytochemical screening of 80% methanol crude extract and solvent fractions of *R. nepalensis*.

| Metabolites | Test | 80%<br>Methanol<br>Extract | Aqueous<br>Fraction | Ethyl<br>acetate<br>Fraction | Hexane<br>Fraction |
| --- | --- | --- | --- | --- | --- |
| Alkaloids | Wagner's Test | + | + | + | - |
| Anthraquinones | Borntrager's Test | + | + | + | + |
| Flavonoids | Shinoda's test | + | + | + | - |
| Glycosides | Keller-Killani test | + | - | + | + |
| Phenols | Lead acetate | + | + | + | - |
| Saponins | Foam Test | + | + | - | - |
| Phytosterols | Salkowski Test | + | - | + | + |
| Tanins | FeCl<sub>3</sub> | + | + | + | - |
| Triterpinoides | Salkowski Test | + | + | + | - |

**Notes:** + = Present, - = Absent

### Acute oral toxicity

Female mice were subjected to an acute oral toxicity test in accordance with the OECD-425 recommendations. The test found that a limit dosage of 2000 mg/kg of *R. nepalensis* hydromethanolic crude extract produced no mortality over the first 24 hours and for the next 15 days. The experimental animals’ physical and behavioral evaluations likewise indicated no obvious indicators of acute toxicity. As a result, the extract’s LD50 value is established to be more than 2000mg/kg.

### Urinary output

The medium dose (M200) of the 80% methanol extract generated a significant elevation in urine production (p<0.001) compared to the negative control at the conclusion of the second, fourth, and fifth hour periods. The highest dose (M400) of the 80% methanol extract generated a significant rise in urine production (p<0.001) compared to the negative control commencing from the second hour until the end of the fifth hour periods of urine collection. Except towards the end of the fifth hour period (p<0.001), the standard drug had a comparable profile in terms of urine production to M400. (See **Table 3**)

**Table 3.** The effect of crude 80% methanol extract of *R. nepalensis* on urine volume in mice.

| Group | Volume of Urine (mL) |  |  |  |  | Diuretic<br>Action | Diuretic<br>Activity |
| --- | --- | --- | --- | --- | --- | --- | --- |
|  | 1 h | 2 h | 3 h | 4 h | 5 h |  |  |
| **2% TW80** | 0.34 ± 0.01 | 0.45 ± 0.01 | 0.63 ± 0.01 | 0.67 ± 0.02 | 0.77 ± 0.01 |  |  |
| **F10** | 0.47 ± 0.02<sup>a2</sup> | 0.66 ± 0.02<sup>a3</sup> | 0.84 ± 0.02<sup>a3</sup> | 0.98 ± 0.03<sup>a3</sup> | 1.28 ± 0.03<sup>a3</sup> | 1.52 | 1.00 |
| **M100** | 0.35 ± 0.01<sup>b2c2</sup> | 0.58 ± 0.01<sup>a1b1</sup> | 0.64 ±<br>0.02<sup>b3c2d3</sup> | 0.72 ± 0.01<sup>b3c2d3</sup> | 0.82 ± 0.01<sup>b3c3d3</sup> | 1.08 | 0.71 |
| **M200** | 0.42 ± 0.03 | 0.62 ± 0.02<sup>a3</sup> | 0.73 ±<br>0.01<sup>a2b2d2</sup> | 0.85 ± 0.02<sup>a3b2</sup> | 0.99 ± 0.02<sup>a3b3</sup> | 1.28 | 0.84 |
| **M400** | 0.43 ± 0.02 | 0.63 ± 0.01<sup>a3</sup> | 0.82 ± 0.01<sup>a3</sup> | 0.91 ± 0.01<sup>a3</sup> | 1.07 ± 0.03<sup>a3b3</sup> | 1.34 | 0.88 |

**Notes:** Each value represents mean ± S.E.M; n=6; a, against 2% TW80; b, against standard (F10); c, against M200; d, against M400; 1: p < 0.05, 2: p < 0.01, 3: p < 0.001; M100: 80% methanol extract 100 mg/kg, M200: 80% methanol extract 200 mg/kg, M400: 80% methanol extract 400 mg/kg, F10: Furosemide 10 mg/kg, 2% TW80: 2% Tween 80.

During the hours of urine collection, the lowest dose of the ethyl acetate fraction (EAF100) and all extract doses of the n-hexane fraction failed to substantially affect urine output in contrast to the negative control. In comparison to the negative control, the medium dose of ethyl acetate fraction (EAF200) increased urine output significantly (p<0.001) from the second to the fifth hour. Beginning with the first hour of the urine collection periods and continuing until the end of the fifth hour, EAF200 exhibited significant increment in urine output than the negative control (p< 0.001). (See **Table 4**)

**Table 4.** Effect of n-hexane and ethyl acetate fractions of *R. nepalensis* on urinary output in mice.

| Group | Volume of urine (mL) |  |  |  |  | Diuretic Action | Diuretic Activity |
| --- | --- | --- | --- | --- | --- | --- | --- |
|  | 1 h | 2 h | 3 h | 4 h | 5 h |  |  |
| **2% TW80** | 0.23 ± 0.03 | 0.35 ± 0.02 | 0.43 ± 0.02 | 0.53 ± 0.02 | 0.60 ± 0.02 |  |  |
| **F10** | 0.57 ± 0.05<sup>a3</sup> | 0.72 ± 0.02<sup>a3</sup> | 0.92 ± 0.02<sup>a3</sup> | 1.10 ± 0.04<sup>a3</sup> | 1.36 ± 0.02<sup>a3</sup> | 2.14 | 1 |
| **HF100** | 0.23 ± 0.02<sup>b3</sup> | 0.33 ± 0.02<sup>b3</sup> | 0.47 ± 0.02<sup>b3</sup> | 0.58 ± 0.02<sup>b3</sup> | 0.63 ± 0.01<sup>b3</sup> | 1.10 | 0.51 |
| **HF200** | 0.27 ± 0.04<sup>b3</sup> | 0.34 ± 0.02<sup>b3</sup> | 0.45 ± 0.02<sup>b3</sup> | 0.57 ± 0.02<sup>b3</sup> | 0.62 ± 0.02<sup>b3</sup> | 1.07 | 0.50 |
| **HF400** | 0.32 ± 0.02<sup>b3</sup> | 0.37 ± 0.04<sup>b3</sup> | 0.49 ± 0.03<sup>b3</sup> | 0.58 ± 0.02<sup>b3</sup> | 0.65 ± 0.02<sup>b3</sup> | 1.13 | 0.53 |
| **EAF100** | 0.27 ± 0.02<sup>b3</sup> | 0.39 ± 0.01<sup>b3e3f3</sup> | 0.49 ± 0.02<sup>b3e3f3</sup> | 0.63 ± 0.01<sup>b3e3f3</sup> | 0.67 ± 0.02<sup>b3e3f3</sup> | 1.14 | 0.53 |
| **EAF200** | 0.33 ± 0.02<sup>b3</sup> | 0.53 ± 0.03<sup>a3b3f3</sup> | 0.75 ± 0.02<sup>a3b3f3</sup> | 0.83 ± 0.03<sup>a3b3f2</sup> | 0.95 ± 0.02<sup>a3b3f3</sup> | 1.64 | 0.77 |
| **EAF400** | 0.38 ± 0.02<sup>a2b3</sup> | 0.67 ± 0.01<sup>a3</sup> | 1.05 ± 0.04<sup>a3b2</sup> | 1.00 ± 0.04<sup>a3</sup> | 1.18 ± 0.03<sup>a3b2</sup> | 2.00 | 0.93 |

**Notes:** Each value represents mean ± S.E.M (n=6) and was analyzed by ANOVA followed by Tuckey's post-hoc multiple comparison test. a, 2% TW80, b, against F10, c, against HF200 mg/kg, d, against HF400 mg/kg, e, against EAF200 mg/kg, f, against EAF400 mg/kg 1: p < 0.05, 2: p < 0.01, 3: p < 0.001. HF100: n-hexane fraction 100 mg/kg, HF200: n-hexane fraction 200 mg/kg, HF400: n-hexane fraction 400 mg/kg, EAF100: ethyl acetate fraction 100 mg/kg, EAF200: ethyl acetate fraction 200 mg/kg, EAF400: ethyl acetate fraction 400 mg/kg, F10: Furosemide 10 mg/kg, 2%TW80: 2% Tween 80

Only at the end of the fifth hour of urine collection did the lowest dose of aqueous fraction (AQF100) significantly increase urine production (p<0.05) as compared to the negative control. The medium dosage of the aqueous fraction (AQF200) considerably enhanced (p<0.001) urine production as compared to the negative control, beginning in the first hour and continuing until the conclusion of the fifth hour period. Similar to AQF200, the highest dose of the aqueous fraction (AQF400) also induced significant increment (p<0.001) in urine output starting from the first hour to the end of the fifth hour period of urine collection. (See **Table 5**)

**Table 5.**
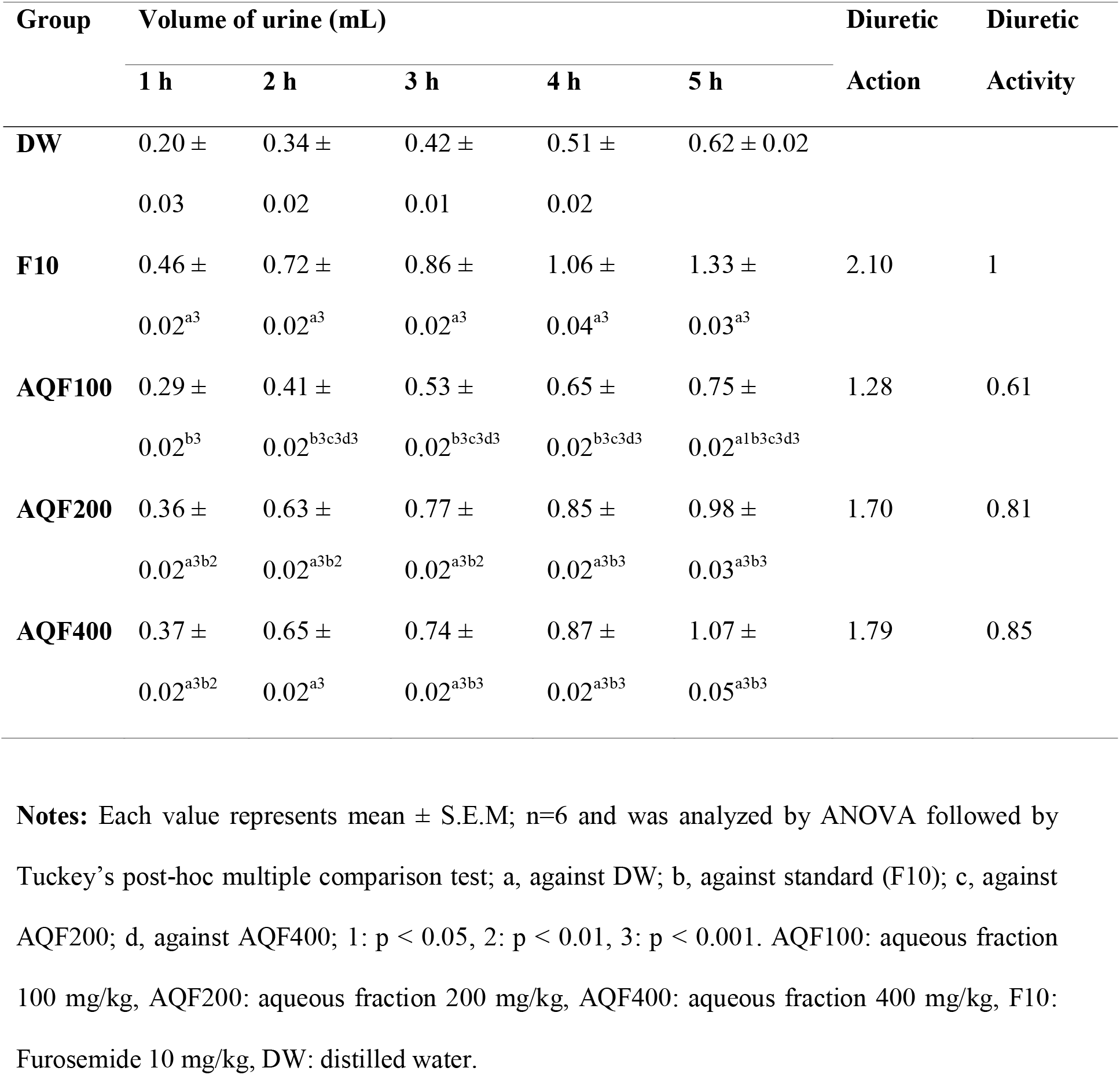
The effect of aqueous fraction of *R. nepalensis* leaves on urine output in mice.

### Urinary electrolyte excretion

When compared to the negative control, the natriuresis induced by the methanol extract at doses M100 (lowest dose), M200, and M400 was considerably greater (p<0.05, p<0.001, and p<0.001 correspondingly; see **Table 6**). The 80% methanol extract doses all caused significant kaliuresis (p<0.001) when compared to the negative control. It was also observed that the excretion of chloride followed a pattern similar to that of potassium, with M100, M200, and M400 producing considerably greater (p<0.001) chloride excretion than the negative control.

**Table 6.**
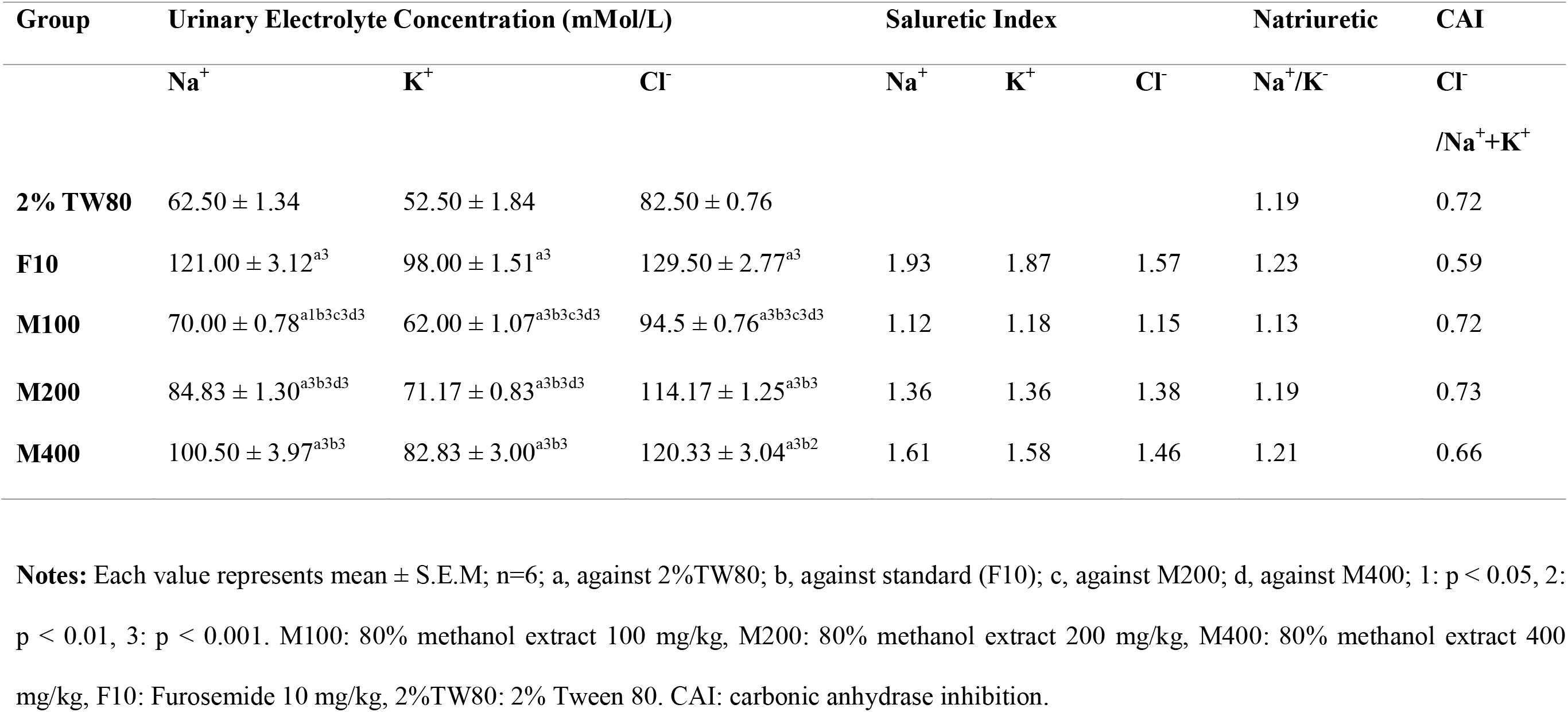
Effect of 80% methanol extract of *R.nepalensis* on urinary electrolyte excretion on mice.

| Group | Urinary Electrolyte Concentration (mMol/L) |  |  | Saluretic Index |  |  | Natriuretic | CAI |
| --- | --- | --- | --- | --- | --- | --- | --- | --- |
|  | Na<sup>+</sup> | K<sup>+</sup> | Cl<sup>-</sup> | Na<sup>+</sup> | K<sup>+</sup> | Cl<sup>-</sup> | Na<sup>+</sup>/K<sup>+</sup> | Cl<sup>-</sup><br>/Na<sup>+</sup>+K<sup>+</sup> |
| **2% TW80** | 62.50 ± 1.34 | 52.50 ± 1.84 | 82.50 ± 0.76 |  |  |  | 1.19 | 0.72 |
| **F10** | 121.00 ± 3.12<sup>a3</sup> | 98.00 ± 1.51<sup>a3</sup> | 129.50 ± 2.77<sup>a3</sup> | 1.93 | 1.87 | 1.57 | 1.23 | 0.59 |
| **M100** | 70.00 ± 0.78<sup>a1b3c3d3</sup> | 62.00 ± 1.07<sup>a3b3c3d3</sup> | 94.5 ± 0.76<sup>a3b3c3d3</sup> | 1.12 | 1.18 | 1.15 | 1.13 | 0.72 |
| **M200** | 84.83 ± 1.30<sup>a3b3d3</sup> | 71.17 ± 0.83<sup>a3b3d3</sup> | 114.17 ± 1.25<sup>a3b3</sup> | 1.36 | 1.36 | 1.38 | 1.19 | 0.73 |
| **M400** | 100.50 ± 3.97<sup>a3b3</sup> | 82.83 ± 3.00<sup>a3b3</sup> | 120.33 ± 3.04<sup>a3b2</sup> | 1.61 | 1.58 | 1.46 | 1.21 | 0.66 |

**Notes:** Each value represents mean ± S.E.M; n=6; a, against 2%TW80; b, against standard (F10); c, against M200; d, against M400; 1: p < 0.05, 2: p < 0.01, 3: p < 0.001. M100: 80% methanol extract 100 mg/kg, M200: 80% methanol extract 200 mg/kg, M400: 80% methanol extract 400 mg/kg, F10: Furosemide 10 mg/kg, 2%TW80: 2% Tween 80. CAI: carbonic anhydrase inhibition.

Significant natriuresis was produced by the highest dose (HF400) of n-hexane fraction in contrast to the negative control (p<0.001). Contrary to the sodium excretion, all the n-hexane extract doses were able to induce significant (p<0.001) potassium and chloride excretion when compared to the negative control. EAF200 and the highest dose (EAF400) of ethyl acetate fraction displayed a higher natriuresis profile compared to the negative control, with a significance level of (p<0.01) and (p<0.001) respectively. A significant potassium excretion effect was noted by EAF200 compared to the negative control. The extract dose of EAF 400, on the other hand, was likewise associated with a greater excretion of potassium and chloride (p<0.001) in comparison to the negative control. (See **Table 7**)

**Table 7.** Effect of n-hexane and ethyl acetate fractions of *R. nepalensis* on urinary electrolyte excretion in mice.

| Group | Urinary Electrolyte Concentration (mMol/L) |  |  | Saluretic Index |  |  | Natriuretic | CAI |
| --- | --- | --- | --- | --- | --- | --- | --- | --- |
|  | Na<sup>+</sup> | K<sup>+</sup> | Cl<sup>-</sup> | Na<sup>+</sup> | K<sup>+</sup> | Cl<sup>-</sup> | Na<sup>+</sup>/K<sup>+</sup> | Cl<sup>-</sup>/Na<sup>+</sup>+K<sup>+</sup> |
| **2% TW80** | 64.83 ± 0.95 | 48.67 ± 0.89 | 95.50 ± 1.77 |  |  |  | 1.33 | 0.84 |
| **F10** | 108.50 ± 1.67<sup>a3</sup> | 91.33 ± 1.56<sup>a3</sup> | 122.17 ± 1.66<sup>a3</sup> | 1.67 | 1.88 | 1.23 | 1.19 | 0.61 |
| **HF100** | 65.50 ± 2.01<sup>b3d3</sup> | 61.00 ± 1.65<sup>a3b3d3</sup> | 108.67 ± 1.52<sup>a3b3</sup> | 1.01 | 1.25 | 1.14 | 1.07 | 0.86 |
| **HF200** | 66.83 ± 0.70<sup>b3d3</sup> | 64.83 ± 1.89<sup>a3b3d1</sup> | 112.33 ± 1.80<sup>a3b2</sup> | 1.03 | 1.33 | 1.18 | 1.03 | 0.85 |
| **HF400** | 78.00 ± 1.16<sup>a3b3</sup> | 70.67 ± 0.88<sup>a3b3</sup> | 110.83 ± 1.68<sup>a3b3</sup> | 1.20 | 1.45 | 1.16 | 1.10 | 0.75 |
| **EAF100** | 65.67 ± 1.05<sup>b3e2f3</sup> | 45.17 ± 1.08<sup>b3f3</sup> | 87.83 ± 1.01<sup>a2b3e2f3</sup> | 1.01 | 0.93 | 0.92 | 1.45 | 0.79 |
| **EAF200** | 71.83 ± 0.91<sup>a2f3</sup> | 55.50 ± 0.43<sup>a3b3</sup> | 96.67 ± 0.88<sup>b3f3</sup> | 1.11 | 1.14 | 1.01 | 1.29 | 0.76 |
| **EAF400** | 97.00 ± 1.03<sup>a3b3</sup> | 59.83 ± 1.01<sup>a3b3</sup> | 106.83 ± 0.87<sup>a3b3</sup> | 1.50 | 1.23 | 1.12 | 1.62 | 0.68 |

**Notes:** Each value represents mean ± S.E.M (n=6) and was analyzed by ANOVA followed by Tuckey's post-hoc multiple comparison test. a, against 2% TW80, b, against F10, c, against HF200 mg/kg, d, against HF400 mg/kg, e, against EAF200 mg/kg, f, against EAF400 mg/kg 1:p < 0.05, 2:p < 0.01, 3:p < 0.001. HF100: n-hexane fraction 100mg/kg, HF200: n-hexane fraction 200mg/kg, HF400: n-hexane fraction 400mg/kg, EAF100: ethyl acetate fraction 100mg/kg, EAF200: ethyl acetate fraction 200mg/kg, EAF400: ethyl acetate fraction 400mg/kg, F10: Furosemide 10 mg/kg, 2%TW80: 2% Tween 80. CAI: carbonic anhydrase inhibition.

AQF200 induced a higher natriuresis effect (p<0.01) compared to the negative control. The potassium and chloride excretion effects of AQF200 were also noticeably higher (p<0.001) compared to the negative control. AQF400 induced significant (p<0.001) excretion of all the measured ions compared to the negative control. As opposed to the negative control, AQF100 caused a substantial (p<0.001) excretion of only chloride ions. (See **Table 8**)

**Table 8.** Effect of the aqueous fraction on urinary electrolyte excretion in mice.

| Group | Urinary Electrolyte Concentration (mMol/L) |  |  | Saluretic Index |  |  | Natriuretic | CAI |
| --- | --- | --- | --- | --- | --- | --- | --- | --- |
|  | Na<sup>+</sup> | K<sup>+</sup> | Cl<sup>-</sup> | Na<sup>+</sup> | K<sup>+</sup> | Cl<sup>-</sup> | Na<sup>+</sup>/K<sup>+</sup> | Cl<sup>-</sup>/Na<sup>+</sup> + K<sup>+</sup> |
| DW | 54.50 ± 9.72 | 45.67 ± 0.67 | 93.83 ± 0.60 |  |  |  | 1.19 | 0.94 |
| F10 | 113.17 ± 1.54<sup>a3</sup> | 96.67 ± 1.05<sup>a3</sup> | 126.50 ± 1.18<sup>a3</sup> | 2.08 | 2.12 | 1.35 | 1.17 | 0.60 |
| AQF100 | 66.83 ± 0.95<sup>b3d1</sup> | 47.67 ± 1.12<sup>b3c3d3</sup> | 109.50 ± 1.18<sup>a3d3</sup> | 1.23 | 1.04 | 1.17 | 1.40 | 0.96 |
| AQF200 | 79.83 ± 1.33<sup>a2b3</sup> | 62.67 ± 0.92<sup>a3b3d3</sup> | 110.50 ± 0.89<sup>a3d3</sup> | 1.46 | 1.37 | 1.18 | 1.27 | 0.78 |
| AQF400 | 87.00 ± 0.97<sup>a3b2</sup> | 78.00 ± 1.15<sup>a3b3</sup> | 118.50 ± 0.76<sup>a3</sup> | 1.60 | 1.71 | 1.26 | 1.12 | 0.72 |

**Notes:** Each value represents mean ± S.E.M; n=6 and was analyzed by ANOVA followed by Tuckey's post-hoc multiple comparison test; a, against DW; b, against standard (F10); c, against AQF200; d, against AQF400; 1: p < 0.05, 2: p < 0.01, 3: p < 0.001; AQF100: aqueous fraction 100 mg/kg, AQF200: aqueous fraction 200 mg/kg, AQF400: aqueous fraction 400 mg/kg, F10: Furosemide 10 mg/kg, DW: distilled water. CAI: carbonic anhydrase inhibition.

### Urinary pH

Urinary measurement of pH demonstrated that the various treatment groups of both the 80% methanol extract and the solvent fractions generated alkaline urine. The pH of urine treated with the 80% methanol extract has demonstrated a rise from M100 (7.09) to M400 (8.05). The control group generated the lowest pH, while the standard group had an intermediate pH (7.78) between the vehicle and extract treated groups. (**See Figure 1**). Treatment with the n-hexane fraction elevated the pH from 7.11 for HF100 to 7.27 for HF400. Similarly, the ethyl acetate fraction raised pH from 7.23 for EAF100 to 8.30 for EAF400. (**See Figure 2**). Similar to the other extracts, treatment with the aqueous fraction elevated pH from 7.54 for AQF100 to 8.24 for AQF400. Treatment with the control generated the lowest pH, and treatment with the standard drug created a pH of 8.01 (**see Figure 3**).

**Figure 1.**
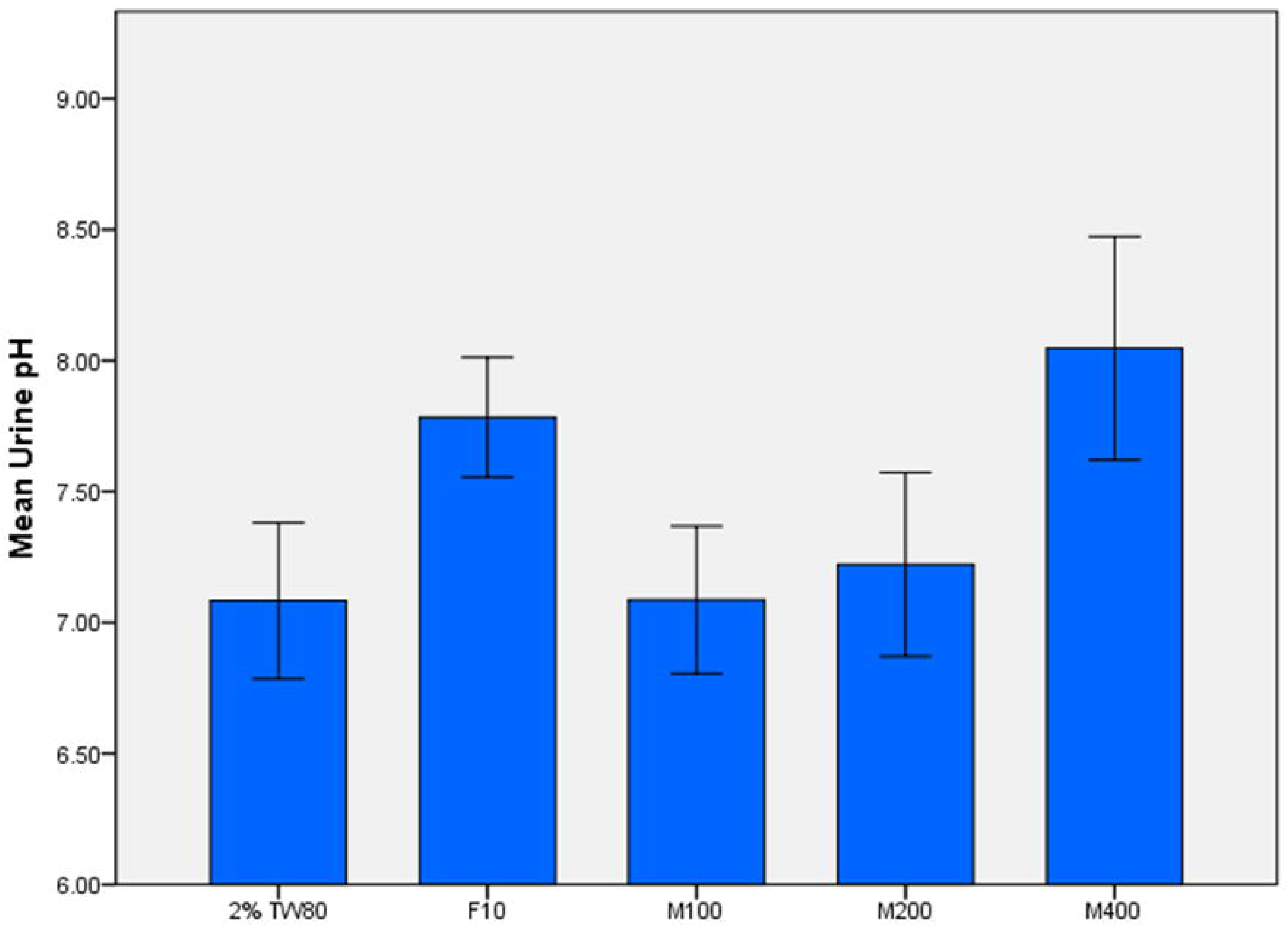
Urinary pH of mice treated with 80% methanol extract of the leaves of *R.nepalensis*. **Notes:** M100: 80% methanol extract 100 mg/kg, M200: 80% methanol extract 200 mg/kg, M400: 80% methanol extract 400 mg/kg, F10: Furosemide 10 mg/kg, 2%TW80: 2% Tween 80.

**Figure 2.**
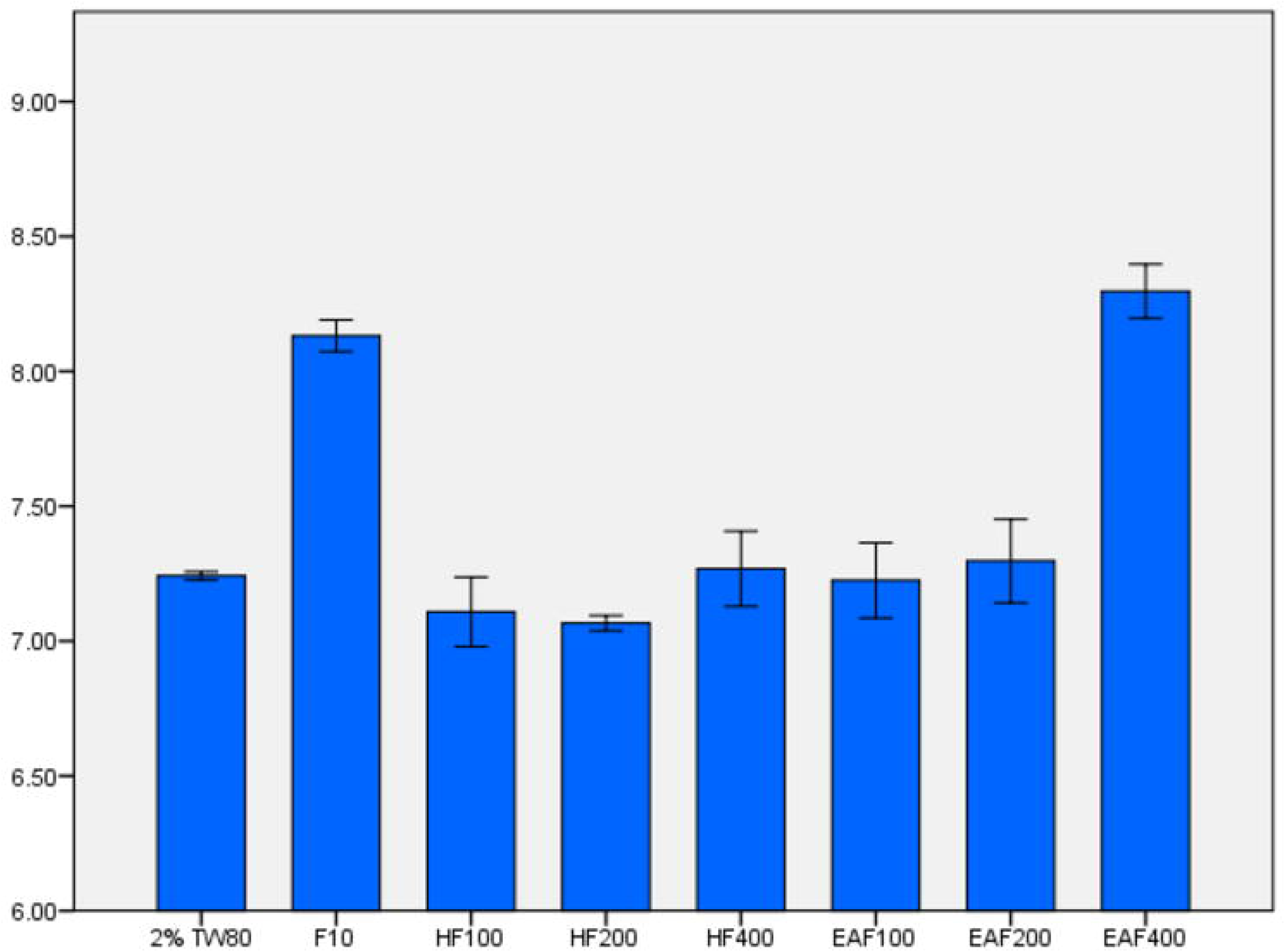
Urinary pH of mice treated with n-hexane and ethyl acetate fractions of the leaves of *R.nepalensis*. **Notes:** HF100: n-hexane fraction 100mg/kg, HF200: n-hexane fraction 200mg/kg, HF400: n-hexane fraction 400mg/kg, EAF100: ethyl acetate fraction 100mg/kg, EAF200: ethyl acetate fraction 200mg/kg, EAF400: ethyl acetate fraction 400mg/kg, F10: Furosemide 10 mg/kg, 2%TW80: 2% Tween 80.

**Figure 3.**
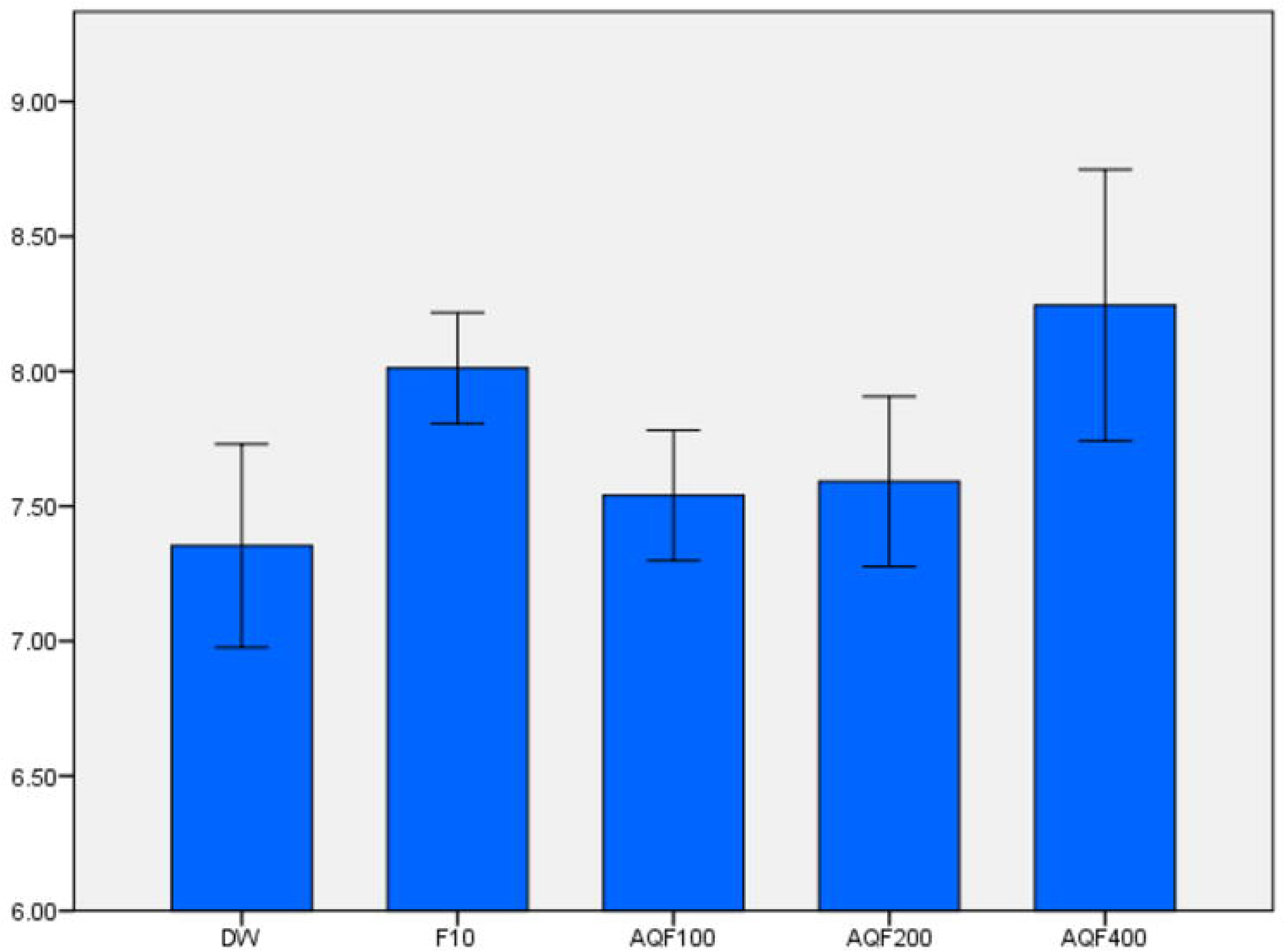
Urinary pH of mice treated with the fractions of the leaves of *R.nepalensis*. **Notes:** AQF100: aqueous fraction 100 mg/kg, AQF200: aqueous fraction 200 mg/kg, AQF400: aqueous fraction 400 mg/kg, F10: Furosemide 10 mg/kg, DW: distilled water.

## Discussion

Diuretics are drugs that help the body eliminate extra water, salt, toxins, and waste products. As a result, they can be employed for the management of various health conditions, mainly heart failure, hypertension, and ascites. (23) The parameters evaluated in this study to examine the diuretic action of the leaf of *R. nepalensis* in mice were urine volume, urine electrolyte, and urinary pH.

M100 failed to substantially enhance urine production over the study period. M200 and M400 were able to considerably promote diuresis. This could reflect the fact that the lowest dose of this extract was below the minimum effective dose and hence did not cause diuresis.(24)

EAF400 and EAF200 were able to significantly cause diuresis. AQF200 and AQF400 also significantly increased diuresis. The lack of any significant urine production by the n-hexane fraction could be due to the elevated concentration of diuretic active metabolites in the methanolic extract, as well as variations in the type, quality, and amount of these active principles present in the three solvent fractions.(25)

Based on the estimated diuretic activity value, the diuretic potential of the 80% methanol crude extract and its solvent fractions was ranked as ‘mild’ and ‘nil’. The 80% methanol crude extract displayed a ‘mild’ type of diuretic activity with a value of 0.84 and 0.88 for M200 and M400, respectively. Similarly, EAF200, EAF400, AQF200, and AQF400 also displayed a ‘mild’ type of diuretic activity, with values of 0.77 and 0.93 for the EAF200 and EAF400 and 0.81 and 0.85 for AQF200 and AQF400, respectively. In contrast to the others, the n-hexane fraction displayed a ‘nil’ diuretic activity at all dose levels with values less than 0.72. Diuretic activity is regarded as good if it exceeds 1.50, moderate if it falls between 1.00 and 1.50, mild if it falls between 0.72-0.99, and zero if it falls below 0.72.(24)

Loop diuretics, like furosemide, increase the excretion of sodium and chloride ions by blocking the Na-K-2Cl cotransporter at the apical membrane of the thick ascending limb of the loop of Henle. (26) Despite the lack of mechanistic study in this research, given the comparable diuretic and saluretic activity of M400, EAF400, and AQF400 to furosemide, it is reasonable to believe that the active components of the crude extract and solvent fractions might have a furosemide-like effect. It is therefore rational to suppose that the extracts of *R. nepalensis* exert their diuretic activity by inhibiting tubular reabsorption of water and accompanying ions. *Rumex abyssinicus Jacq*, another plant in the same genus and family as *R. nepalensis*, has already been recommended for similar activity.(27)

M400, EAF400, and AQF400, on the other hand, increased urinary sodium, potassium, and chloride excretion with significant urine alkalinization, implying that the diuretic activity of these extracts may differ from loop diuretics because loop diuretics increase sodium ion loading to the distal tubule of the nephron, which acidifies the urine by activating H^+^ secretion via the H^+^-ATPase in α intercalated cells.(28) As a result, it is acceptable to conclude that the metabolites found in the leaves of *R. nepalensis* seem to have a distinct diuretic mode of action.

Potassium excess, which occurs when the nephron tubules are unable to absorb it, leads in osmotic type of urine excretion.(27) Quantitative determination of the ions present in the 80% methanol extract and solvent fractions of *R. nepalensis* revealed undetectable amounts of potassium salts. This suggests that the diuretic effect produced by the methanol extract, ethyl acetate fraction, and aqueous fraction does not seem to be of the osmotic type.

A satisfactory natriuresis, a favorable natriuresis, and a favorable K+-sparing activity, respectively, are indicated by Na^+^/K^+^ ratios of >1, 2, and 10.(25, 29) Accordingly, the medium and highest doses of the 80% methanol crude extract displayed a satisfactory natriuresis with an index value of 1.19 and 1.21 for M200 and M400, respectively. Similarly, EAF200 and EAF400 displayed satisfactory natriuresis with values of 1.29 and 1.62, respectively. A similar pattern of natriuresis was also noted for the aqueous fraction, where AQF200 and AQF400 displayed satisfactory natriuresis with values of 1.27 and 1.12, respectively.

Moreover, the crude extract and fractions did not show any reduction in urinary potassium excretion levels, unlike some plant extracts that have been reported to have a potassium sparing effect, like in the case of 1000 mg/kg aqueous extract dose of *Ajuga remota Benth* leaves (20), demonstrating the plant’s ineffectiveness as a potassium-sparing diuretic. This would signal that the extract was ineffective at preventing hypokalemia, which is a common adverse effect of loop diuretic drugs.

The effect of a test substance on the Cl^-^/Na^+^+K^+^ ratio can predict the action of the substance on carbonic anhydrase enzyme activity(CAI).(25) The CAI effect is disregarded if the Cl^-^/Na^+^+K^+^ ratio is between 0.8 and 1.00, but if it is less than 0.8, it is considered to have strong CAI activity.(30) In this study, both M400 and EAF400 exhibited a strong CAI effect with a value of 0.66 and 0.68, respectively. This suggests that M400 and EAF400 might possibly have an inhibiting effect on the renal tubules’ carbonic anhydrase enzyme. This is further supported by the greatly raised urine pH values associated with these specific extract doses, which were 8.05 and 8.30, respectively.

The 80% methanol extract and solvent fractions underwent preliminary phytochemical analysis, which identified many secondary metabolites that seemed to be distributed differentially across the crude extract and solvent fractions. These secondary metabolites have been investigated in the past and are known to have many diuretic effects. Alkaloids, for instance, reduce the activity of carbonic anhydrase enzyme, boost renal blood flow by widening renal afferent arteries, and restrict the reabsorption of water and electrolytes in the tubules.(31) Flavonoids have a diuretic impact as well as a potassium-saving function.(32–34) Phenols also have an inhibiting effect on carbonic anhydrase.(24) Therefore, it is conceivable that the alkaloids, flavonoids, and phenols included in the *R. nepalensis* 80% methanol extract, ethyl acetate, and aqueous fractions may be responsible for the diuretic effects seen in these extracts.

## Conclusion

The findings of this study suggested that the leaf extracts of *R. nepalensis* have a diuretic effect, as evidenced by increased salt and water excretion. This finding substantiate the long-standing folkloric use of *R. nepalensis* leaves as a diuretic agent.

## Acknowledgments

I would like to thank Mr. Abreham Degu, Mr. Alemante Tafesse, and Mr. Zemenu Wube for their unreserved assistance during the course of laboratory work. I would also like to thank Mr. Getenet Chekole for his kind help at the time of plant identification and authentication.

